# Developmental emergence of sleep rhythms enables long-term memory capabilities in *Drosophila*

**DOI:** 10.1101/2022.02.03.479025

**Authors:** Amy R. Poe, Lucy Zhu, Patrick D. McClanahan, Milan Szuperak, Ron C. Anafi, Andreas S. Thum, Daniel J. Cavanaugh, Matthew S. Kayser

## Abstract

In adulthood, sleep-wake rhythms are one of the most prominent behaviors under circadian control. However, during early life, sleep is spread across the 24-hour day (1–4). The mechanism through which sleep rhythms emerge, and the consequent advantage conferred to a juvenile animal, are unknown. In 2^nd^ instar *Drosophila* larvae (L2), like human infants, sleep is not under circadian control (5). Here, we identify the precise developmental timepoint when the circadian clock begins to regulate sleep in *Drosophila*, leading to the emergence of sleep rhythms at the early 3^rd^ instar stage (L3). At this stage, a cellular connection forms between DN1a clock neurons and arousal-promoting Dh44 neurons, bringing arousal under clock control to drive the emergence of circadian sleep. Finally, we demonstrate that L3 but not L2 larvae exhibit long-term memory (LTM) of an aversive cue, and that this LTM depends upon deep sleep generated once sleep rhythms begin. We propose that the developmental emergence of circadian sleep enables more complex cognitive processes, including the onset of enduring memories.

## Results

### Rhythmic sleep emerges in early 3^rd^ instar larvae

Across species, from flies to mammals, central molecular clocks begin cycling during very early developmental periods (6–10), suggesting that maturation events downstream of the core clock are required to generate rhythmic sleep output. Despite the presence of a functioning circadian clock network (7, 11, 12), sleep in 2^nd^ instar larvae [48 hrs after egg laying (AEL)] is not under clock control (5). To determine when sleep becomes influenced by the circadian system, we examined sleep under constant conditions at 4 times across the day (Circadian Time [CT] 1, CT7, CT13, and CT19) in developmentally age-matched 2^nd^ (L2) and early 3^rd^ instar (L3; 72h AEL) larvae. As expected, there were no differences in sleep across the day in L2 (Fig. 1a-b). However, we observed emergence of a diurnal sleep pattern in L3, specifically an increase in sleep duration driven by more sleep bouts at CT13 and CT19 (subjective dark times) compared to CT1 and CT7 (Fig. 1c-d; Fig. S1a). We next pooled the total sleep bouts for L2 and L3 larvae at CT1 and CT13 and plotted the cumulative probability distribution of sleep bout length. We found that L2 larvae have similar bout length distributions at CT1 and CT13 (Fig. 1e), in contrast to early L3 larvae which exhibit a higher proportion of longer sleep bouts at CT13 compared to CT1 (Fig. 1f) (p>0.5, L2; p<0.0001, L3; Kolmogorov-Smirnov test). To determine if the diurnal pattern of sleep in early L3 is clock-dependent, we examined sleep in mutants for the core clock genes, *per* and *tim* (13, 14). In both *per^0^* and *tim^0^* mutants, we observed no differences across the day in sleep (Fig. 1g-h; Fig. S1b-e); moreover, sleep differences across the day were absent in L3 raised in constant light (LL), which disrupts the molecular clock (Fig. S1f-g). Together, our data define a precise developmental window in which sleep comes under clock control.

**Fig. 1:**
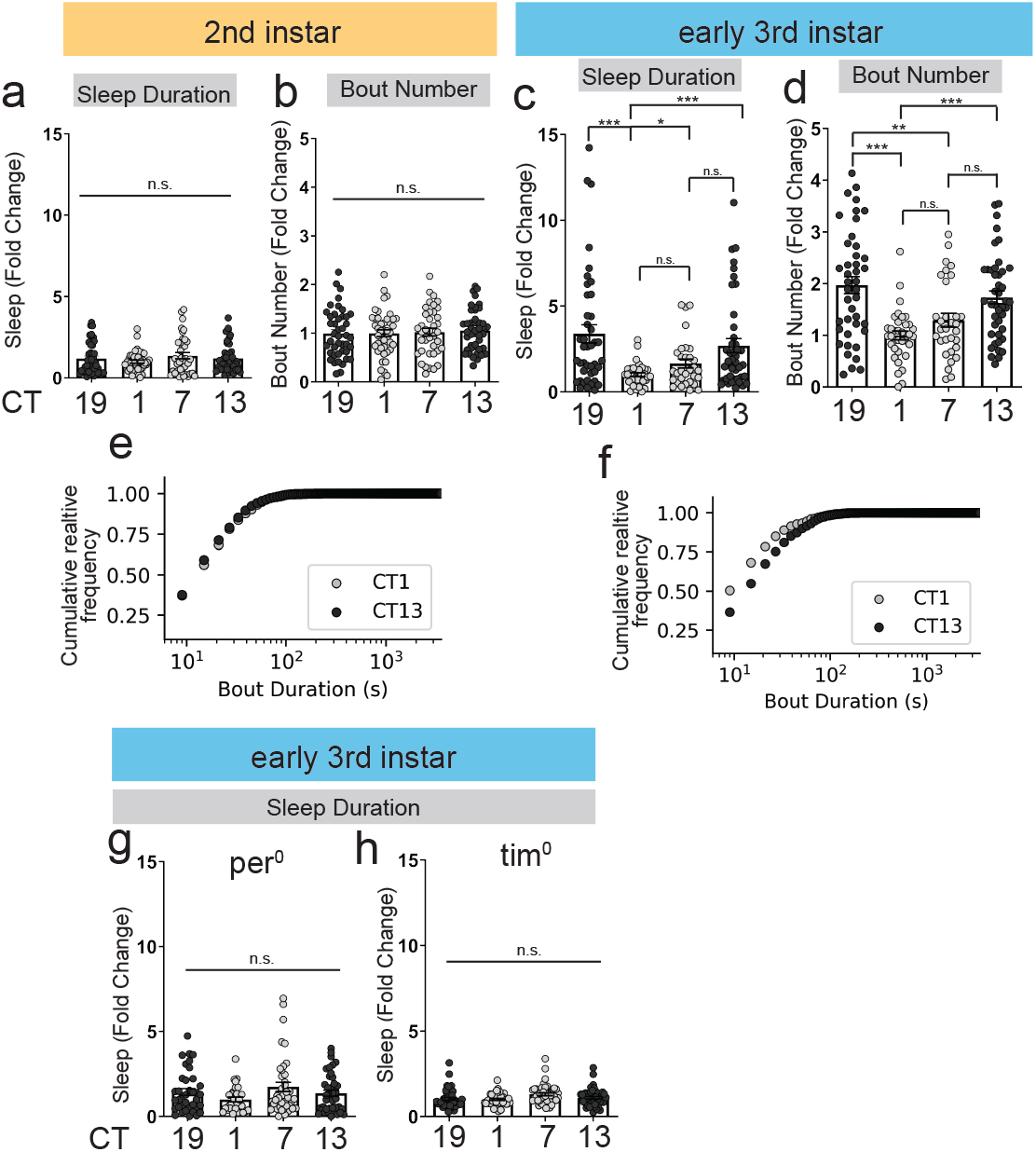
Rhythmic sleep emerges in early L3. **a-b**, L2 Sleep duration and sleep bout number across the day under constant dark conditions. **c-d**, L3 sleep duration and bout number. **e-f**, Cumulative relative frequency of sleep bout duration at CT1 and CT13 for L2 (e) and L3 (f). **g-h**, Sleep duration in L3 clock mutants. Subjective day is shown in light gray circles, subjective night in black. Sleep metrics represent fold change (normalized to avg. value at CT1 for a given panel). a-d, g-h, n=31-44 larvae. One-way ANOVA followed by Tukey’s multiple comparison test [(a-d) and (g-h)]; Kolmogorov-Smirnov test [(e-f)]. n.s., not significant, \**P*<0.05, \*\**P*<0.01, \*\*\**P*<0.001 for this and all subsequent figures.

Although L2 sleep is clock-independent, we previously described developmental changes to sleep duration across this larval stage (5). To determine whether L3 sleep likewise undergoes ontogenetic changes, we held circadian time constant at either a subjective morning (CT5) or subjective evening (CT21) time point and examined sleep duration in newly molted 3^rd^ instar larvae (72h AEL; +0h), larvae aged 4 hours more (76h AEL; +4h), and larvae aged 8 hours (80h AEL; +8h). We did not detect sleep differences in larvae from CT5+0h to CT5+4h, but did find a reduction in sleep by CT5+8h sleep (Fig. S1h); a similar pattern was evident at CT21 (Fig. S1i), suggesting changes to sleep with development occur during L3, perhaps in anticipation of pupation. Under natural conditions, larvae continue to develop as circadian time progresses. Is circadian control of sleep relevant in the face of these developmental changes? We allowed for simultaneous influence of age and circadian time by focusing on L3 sleep patterns at day-night transitions. L3 exhibited a reduction in sleep duration in the night-to-day transition from CT21 to CT1, and an increase in sleep in the day-to-night transition from CT9 to CT13 (Fig. S1j-l). Thus, sleep continues to cycle in relation to circadian time even as larvae develop.

### Dh44 neurons in central brain control arousal in 2^nd^ instar larvae

To investigate potential signaling molecules that regulate the emergence of sleep-wake rhythms, we conducted a neural activation screen of peptidergic neurons. We found that activation of *Dh44-Gal4* expressing neurons using the heat-sensitive cation channel, TrpA1 (15), caused a significant reduction in sleep duration in 2^nd^ instar larvae with little effect on overall activity (Fig. S2a). Acute activation of *Dh44-Gal4* expressing neurons resulted in a loss of sleep primarily due to a reduction in sleep bout length (Fig. 2a-c; Fig. S2b-d). The decrease in sleep was not due to an increase in feeding, as there was no difference in the number of mouth hook contractions normalized to time awake (Fig. S2e). *Dh44*-Gal4 showed restricted expression in the 2^nd^ instar larval brain, labeling only a few cells in the central brain and ventral nerve cord (VNC) (Fig. 2d). To determine if these neurons bidirectionally modulate larval sleep, we expressed the inwardly rectifying potassium channel, Kir2.1 (16), in Dh44 neurons, which led to increased sleep duration, bout length, and bout number (Fig. S2f-h). Additionally, pan-neuronal knockdown of Dh44 increased sleep duration, bout length, and bout number (Fig. 2e-g; Fig. S2i,j). Together, our data indicate that Dh44 neurons bidirectionally modulate arousal in 2^nd^ instar larvae, and that Dh44 is required for maintaining normal arousal at this stage.

**Fig. 2:**
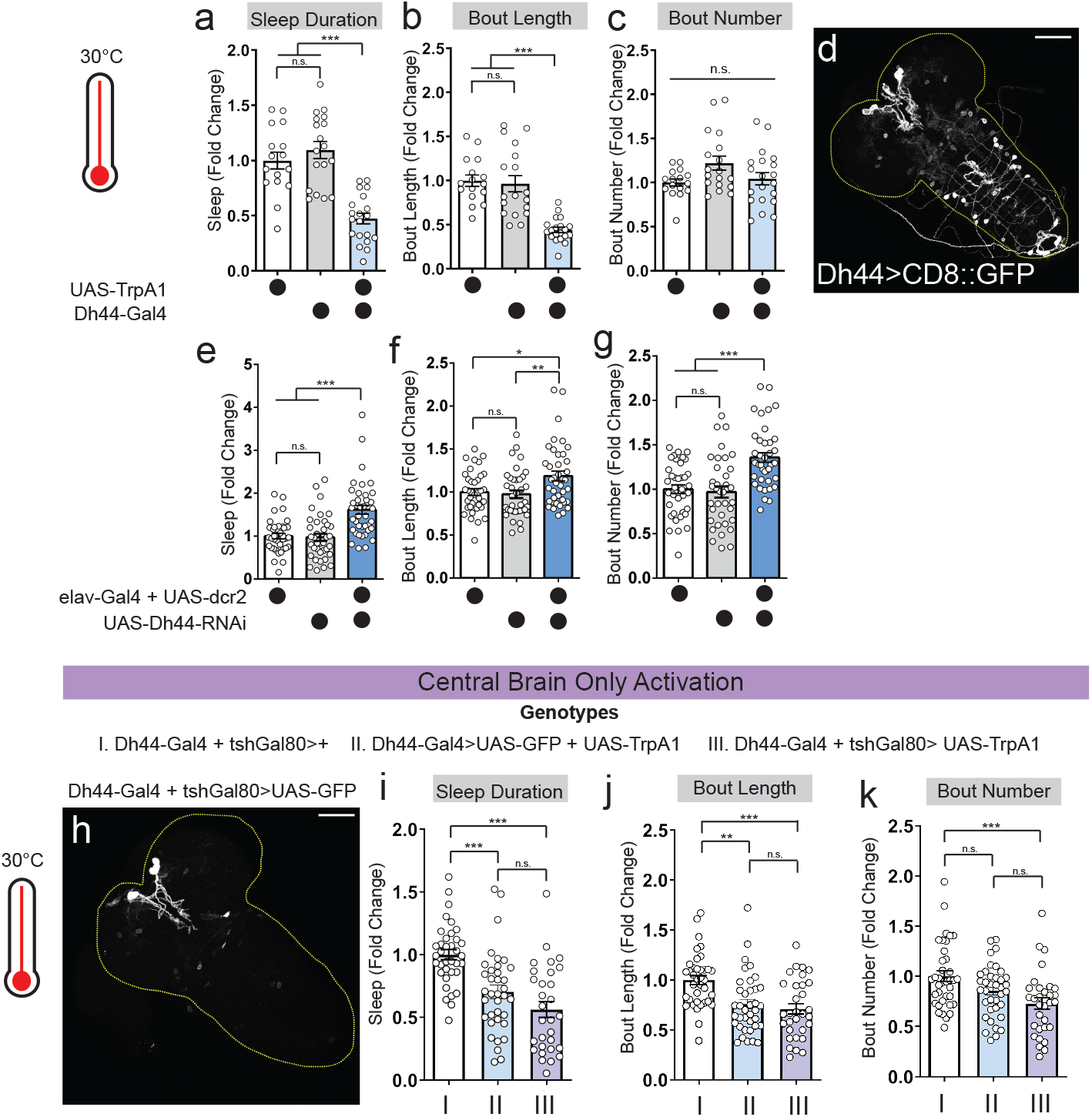
Dh44 neurons in central brain control arousal in L2. **a-c**, Sleep duration, bout length, and bout number in L2 expressing *UAS-TrpA1* with *Dh44-Gal4* and genetic controls. **d**, L2 brain and ventral nerve cord showing GFP expression in Dh44 neurons (*Dh44-Gal4*>*UAS-CD8::GFP*). **e-g**, Sleep duration, bout length, and bout number in L2 expressing *UAS-Dh44-RNAi* with *elav-Gal4* + *UAS-dcr2* and genetic controls. **h**, L2 brain and ventral nerve cord showing GFP expression in larvae expressing *Dh44*-Gal4 in presence of *tshGal80* and *UAS-GFP*. **i-k**, Sleep duration, bout length, and bout number in L2 expressing *Dh44*-Gal4 in presence of *tshGal80* and *UAS-TrpA1* with genetic controls. Sleep metrics represent fold change (normalized to avg. value of control) in this and subsequent figures. a-c, n=16-20 larvae; e-g, n=37-40 larvae; i-k, n=30-36 larvae. One-way ANOVA followed by Tukey’s multiple comparison test. Scale bars=50 microns.

We next asked which subpopulation of *Dh44* neurons is sufficient to induce waking. We used intersectional genetics approaches to restrict *Dh44-Gal4* expression to either the central brain or VNC. Activation of Dh44 neurons only in the central brain decreased sleep duration, bout length, and bout number to a similar extent as activation of all Dh44 cells (Fig. 2h-k; Fig. S3a-c). In contrast, activation of Dh44 neurons in the VNC had no effect on sleep (Fig. S3d-j). Thus, Dh44 neurons in the L2 brain promote arousal.

### Dh44 neurons anatomically & functionally connect with DN1a clock neurons in early L3

Dh44 neurons are part of the *pars intercerebralis* (PI), which in the adult is a circadian output center that conveys timing information from the central clock to downstream effectors (17, 18); additionally, in adulthood, Dh44 neural activity cycles across the day in a clock-dependent fashion (19, 20). We wondered whether Dh44 neurons undergo a functional change during development, taking on a circadian role in addition to promoting arousal. To determine if changes in Dh44 neural activity across the day correlate with the emergence of sleep rhythms in L3, we performed live imaging of larval Dh44 neurons expressing *UAS-GCaMP7f* at CT1 and CT13. While we observed no differences in activity in L2 at CT1 and CT13 (Fig. 3a-c), by early L3 Dh44 activity was lower at CT13 compared to CT11 (Fig. 3d-f), suggesting that Dh44 neurons begin to receive clock input at this stage. To test whether larval clock neurons anatomically connect to Dh44 neurons in early L3, we next assessed synaptic connections between different populations of clock neurons and Dh44 neurons using neurexin-based GFP reconstitution (GRASP) (21, 22). At the L2 and L3 stages, the functional larval clock network is comprised of 4 s-LNvs, 2 DN1as, and 2 DN2s per hemisphere (23–25). Using *cry-Gal4* (larval s-LNv/DN1a-specific driver) with *Dh44-LexA*, we observed distinct puncta around the cell body and dendritic area of Dh44 neurons in early L3 but not L2 (Fig. 3g-j). We did not observe GRASP signal with *pdf-Gal4* (larval s-LNv-specific driver) or *per-Gal4* (larval DN2-specific driver) (Fig. S4a-b). To determine if DN1as and Dh44 neurons are functionally connected in early L3, we expressed ATP-gated P2X2 receptors (26) in DN1a neurons and GCaMP6 in Dh44 neurons. When DN1as were activated with ATP application to dissected brains, we observed an increase in calcium in Dh44 neurons in L3, but not in L2 (Fig. 3k,l). Dh44 neurons, therefore, form anatomical and functional connections with DN1a clock neurons in early 3^rd^ instar larvae, but not prior to this time. Given the connectivity changes upstream of Dh44 neurons between L2 and L3, we next asked whether the functional consequences of Dh44 neuronal activation likewise change. We found that, as in L2, activation of Dh44 neurons in L3 promotes arousal regardless of time of day (Fig. 3m), supporting the hypothesis that upstream input from the clock is the major developmental switch. Thus, Dh44 neurons become connected to the central clock in early L3, bringing arousal under clock control to drive sleep rhythms.

**Fig. 3:**
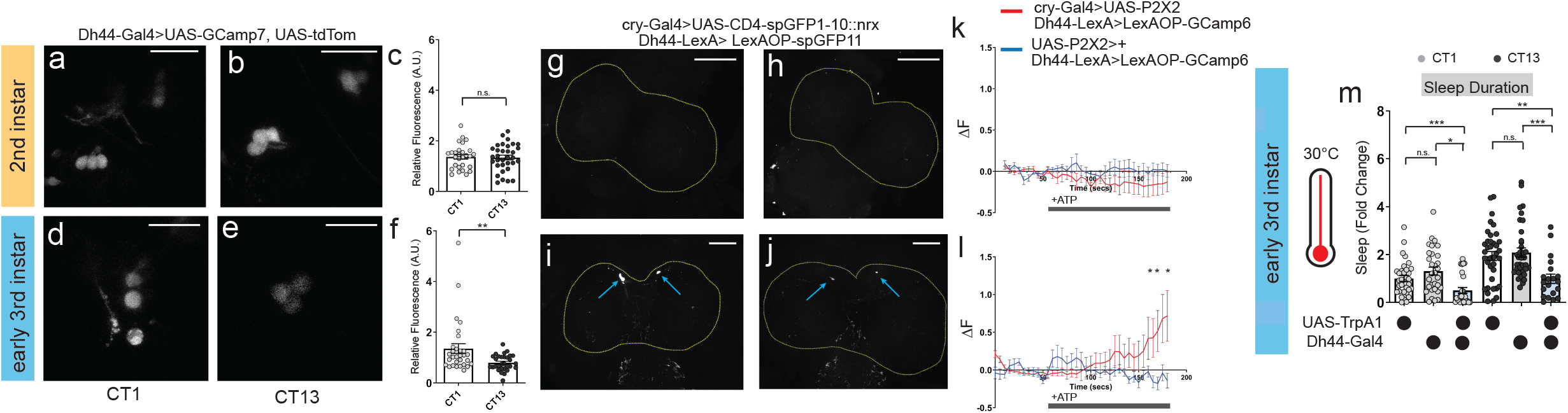
Dh44 neurons anatomically and functionally connect with DN1as in early L3. **a-b**, *Dh44-Gal4*>*UAS-GCaMP7f* expression in L2 brains at CT1 and CT13. **c**, Normalized GCaMP intensity in Dh44 neurons in L2 at CT1 and CT13. n=32-37 cells, 10-11 brains. **d-e**, *Dh44-Gal4*>*UAS-GCaMP7f* expression in L3 brains at CT1 and CT13. **f**, Normalized GCaMP intensity in Dh44 neurons in L3 at CT1 and CT13. n=30 cells, 10 brains. **g-j**, Neurexin-based GFP reconstitution (GRASP) between DN1as (*cry*-Gal4) and Dh44 neurons (*Dh44*-LexA) in L2 (g-h) and L3 (i-j) brains. Yellow dotted lines=central brain; blue arrows=Dh44 cell bodies. n=8-10 brains. **k-l**, GCaMP6 signal in Dh44 neurons with activation of DN1a neurons in L2 (k) and L3 (l). n=9-15 cells, 8 brains. Black bar indicates ATP application. **m**, Sleep duration in L3 expressing *Dh44*-Gal4> *UAS-TrpA1* and genetic controls at CT1 and CT13. n=20-36 larvae. Unpaired two-tailed Student’s *t*-test [(c) and (f)]; Mann-Whitney test [(k) and (l)]; one-way ANOVAs followed by Tukey’s multiple comparison tests (m). Scale bars=25 microns for a-b, c-d. Scale bars=50 microns for g-i.

### CCHa1 signaling between DN1as and Dh44 neurons is necessary for sleep rhythms

We next sought to determine the molecular signals used in the DN1a-Dh44 circuit to drive L3 sleep rhythms. We conducted an RNAi-based candidate screen of receptors for 5 neuropeptides shown to cycle in adult DN1a neurons (27). Sleep duration at CT1 and CT13 was assessed with receptor knockdown in Dh44 neurons, and we found that knockdown of only *CCHamide-1 receptor* (*CCHa1-R*) resulted in a loss of rhythmic changes in sleep duration and bout number (Fig. 4a-b; Fig. S4c). To determine if *CCHamide-1* release from DN1a neurons is required, we knocked down *CCHa1* in DN1as and observed a loss of rhythmic changes in sleep at CT1 and CT13 (Fig. 4c; Fig. S4d). Manipulations of *CCHamide-1* signaling in 2^nd^ instar larvae did not affect sleep duration or bout number at CT1 or CT13 (Fig. 4d-e; Fig. S4e-f), providing further evidence that this circuit is not yet functional at this stage.

**Fig. 4:**
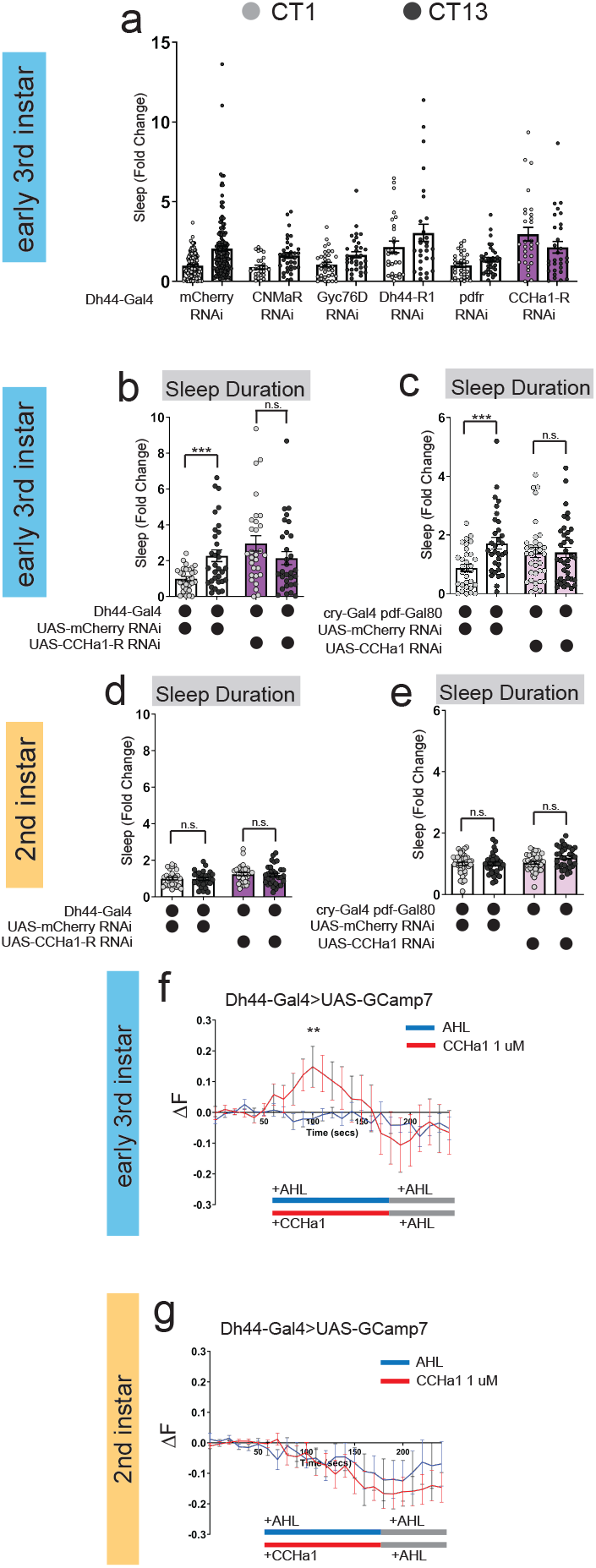
CCHa1 signaling between DN1a and Dh44 neurons is necessary for sleep rhythms. **a**, Sleep duration in L3 at CT1 and CT13 with *Dh44-Gal4*>*UAS-RNAi* targeting neuropeptide receptors. **b-c**, Sleep duration in L3 expressing *UAS-CCHa1-R-RNAi* with *Dh44-Gal4* (b) or *UAS-CCHa1-RNAi* with *cry-Gal4 pdf-Gal80* (DN1as) (c) and genetic controls at CT1 and CT13. **d-e**, Sleep duration in L2 expressing *UAS-CCHa1-R-RNAi* with *Dh44-Gal4* (d) or *UAS-CCHa1-RNAi* with *cry-Gal4 pdf-Gal80* (DN1as) (e) and genetic controls at CT1 and CT13. **f-g**, GCaMP7 signal in Dh44 neurons during bath application of 1 uM CCHa1 synthetic peptide in L3 (f) and L2 (g) brains. n=11-15 cells, 5-7 brains. Red/blue bar indicates timing of CCHa1 (red) or AHL (blue) application and gray bar indicates timing of washout AHL application. Sleep metrics represent fold change (normalized to avg. value of control). a, n=137-144 *mCherry RNAi* larvae, n=27-35 RNAi larvae; b-e, n=29-38 larvae. b-e, Ordinary one-way ANOVAs followed by Sidak’s multiple comparisons tests [(b-e)]; Mann-Whitney test [(f) and (g)].

To confirm that *CCHamide-1* activates Dh44 neurons, we bath applied CCHamide-1 peptide to dissected larval brains expressing *UAS-GCaMP7* and indeed observed an increase in intracellular calcium in Dh44 neurons in early L3 (Fig. 4f). CCHa1 application did not alter calcium levels in Dh44 neurons in L2 (Fig. 4g), indicating that Dh44 neurons are not competent to receive CCHa1 input prior to early L3. Together, these findings demonstrate that a developmentally-timed circuit connecting DN1a and Dh44 neurons utilizes CCHa1 neuropeptidergic signaling to drive circadian sleep.

### Onset of sleep rhythmicity facilitates long-term memory

What advantages are conferred to an animal with the onset of circadian sleep patterns? Sleep is a critical regulator of memory formation and consolidation during development (28, 29). We first asked if sleep rhythms are necessary for short-term memory (STM). We utilized a two-odor reciprocal olfactory conditioning paradigm and tested aversive memory performance in both L2 and L3 immediately after training (30–34). Naïve preferences for odor and aversive stimuli were indistinguishable between L2 and L3 for each genetic manipulation (Fig. S5a-c). We found that both L2 and L3 control animals show STM with similar performance indices (Fig. 5a), indicating that the presence of sleep rhythms is not a prerequisite. Knockdown of CCHa1-R in Dh44 neurons had no effect on STM in early L3, providing further evidence that circadian control of sleep is not required (Fig. 5a).

**Fig. 5:**
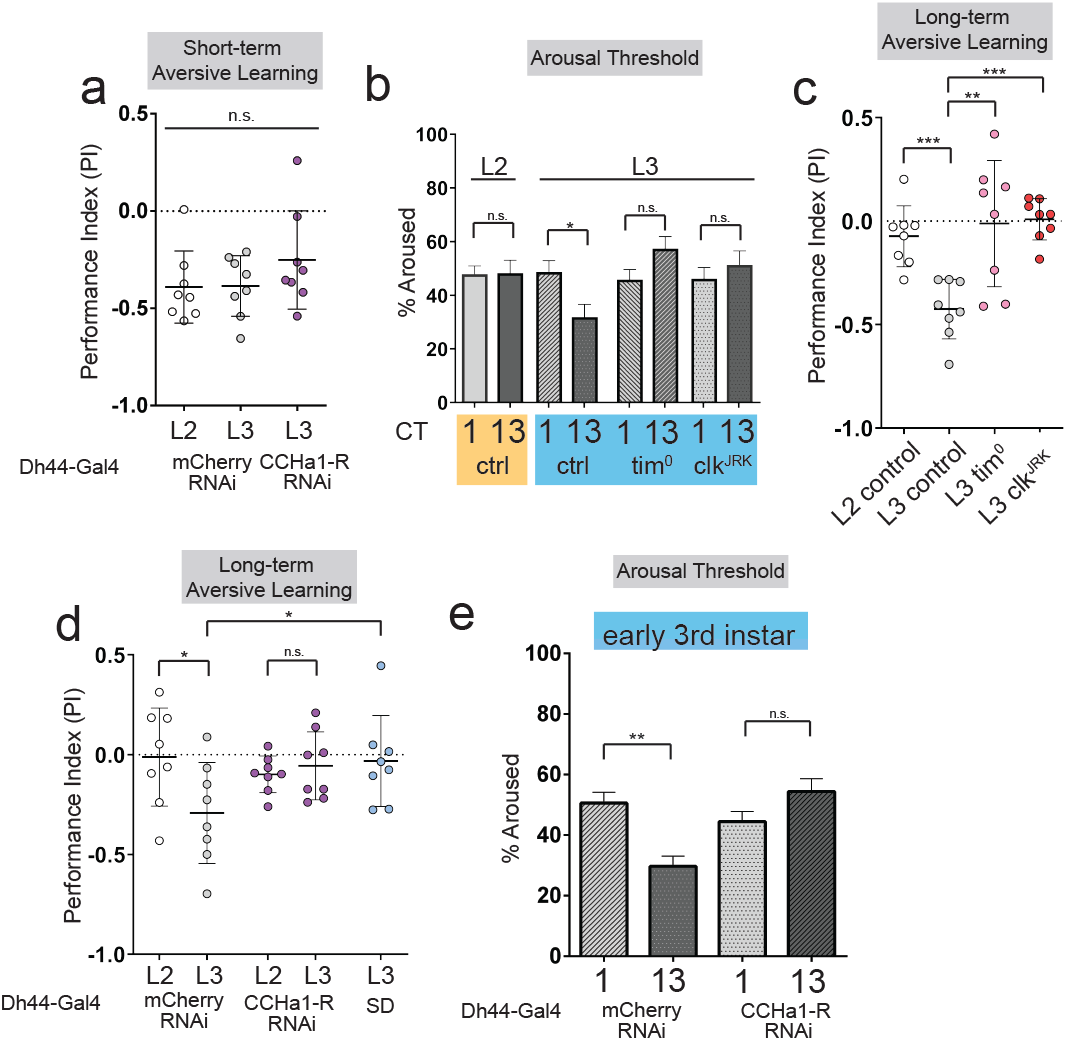
Sleep rhythms facilitate long-term memory formation. **a**, Short-term aversive memory performance in L2 and L3 genetic control (*Dh44-Gal4*>*mCherry RNAi*) and L3 expressing *Dh44-Gal4*>*CCHa1-R-RNAi*. **b**, Arousal threshold in L2 controls (yellow shaded), L3 controls, and L3 *tim^0^* and *clk^JRK^* mutants (blue shaded) at CT1 and CT13 (Per condition: n=230-350 sleep episodes; 36 larvae). **c**, Long-term aversive memory performance in L2 and L3 controls and L3 *tim^0^* and *clk^JRK^* mutants. **d**, Long-term aversive memory performance in L2 and L3 genetic control (*Dh44-Gal4*>*mCherry RNAi*); L2 and L3 expressing *Dh44-Gal4*>*CCHa1-R-RNAi;* and in sleep deprived (SD) L3 genetic controls (*Dh44*-Gal4>*mCherry RNAi*). **e**, Arousal threshold in L3 genetic control (*Dh44-Gal4*>*mCherry RNAi*) and L3 expressing *Dh44*-Gal4>*CCHa1-R-RNAi* at CT1 and CT13 (n=270-430 sleep episodes; 36 larvae). a, c, d, n=8 PIs (240 larvae) per genotype. One-way ANOVA followed by Tukey’s multiple comparison test [(a), (c), and (d)] and ordinary one-way ANOVAs followed by Sidak’s multiple comparisons tests [(b) and (e)].

Deep sleep stages are important for long-term memory (LTM) consolidation across species (35–37). While STM is similar in L2 and L3, we wondered if developmental sleep differences might be relevant for more enduring memories. We therefore asked if clock control of sleep modulates specific sleep features, such as sleep depth, that are beneficial to memory. To assess sleep depth, we examined arousal threshold in sleeping L2 and L3 animals at CT1 or CT13. While L2 showed similar levels of responsiveness to a low intensity light stimulus irrespective of time of day, sleeping L3 were less responsive during the subjective night (Fig. 5b), consistent with deeper sleep during this time. Providing further evidence of increased sleep depth in L3 during the night, L3 exhibited increased probability of transitioning from a wake state to sleep at CT13 compared to CT1, and decreased likelihood of exiting a sleep state; L2 showed no differences in these measures across the day (38) (Fig. S5d-e). Additionally, we observed that deeper sleep in L3 at CT13 was absent in clock mutants (Fig. 5b). These data demonstrate that in addition to an increase in sleep duration at night, L3 also sleep more deeply in a clock-dependent manner.

To test whether circadian induction of deeper sleep in L3 provides advantages to memory function, we assessed aversive LTM based on three odor-quinine training cycles (30, 39). Remarkably, we found that L2 did not show LTM, but early L3 exhibited a strong persistent memory of the aversive cue (Fig. 5c). We then tested if LTM in early L3 is clock-dependent by examining memory performance in *tim^0^* and *clk^JRK^* mutants (40), and found that both mutations rendered L3 unable to show LTM (Fig. 5c). Critically, L3 and L2 *tim^0^* and *clk^JRK^* mutants still show STM (Fig. S5f,j), arguing against a more generalized inability to exhibit memories in clock mutants; moreover, we detected no differences in baseline preference to odors or quinine with regard to larval stage or clock mutants (Fig. S5g-i, k-m). We next examined whether knockdown of *CCHa1-R* in Dh44 neurons, which disrupts sleep rhythms in L3 with no effect in L2, alters LTM. L2 RNAi control larvae did not show LTM (as expected) and *CCHa1-R* knockdown had no further effect; by contrast in L3, knockdown of *CCHa1-R* in Dh44 neurons blocked LTM (Fig. 5d). Consistent with deeper sleep being necessary for LTM, L3 *Dh44*>*CCHa1-R RNAi* larvae failed to exhibit the increased arousal threshold normally observed during the night (Fig. 5e). Finally, we tested if sleep itself is necessary for LTM in early L3 by using a high intensity light stimulus to sleep deprive larvae during the 1.5 hours following training (Fig. S5n). We found that sleep deprivation between training and testing disrupted LTM (Fig. 5d). Our data indicate that sleep rhythms emerge in early 3^rd^ instar larvae to promote deeper sleep at night, which facilitates LTM.

## Discussion

The core molecular clock begins cycling during nascent developmental stages, but numerous behavioral rhythms, including sleep-wake cycles, are not apparent until far later. Here, we discover in *Drosophila* that precisely-timed completion of a circuit motif allows information flow from the central clock network to behavioral output cells, generating sleep rhythms. Our findings suggest that unchecked arousal is a primordial driver of stochastic state changes between sleep and wake during early life, until clock input brings arousal under circadian control. The key molecular signal in this bridge, CCHa1, is known to be involved in circadian intercellular coordination (41), and our results demonstrate an additional clock network-extrinsic role for CCHa1/CCHa1-R in behavioral rhythms. Interestingly, the mammalian homolog of CCHa1-R, gastrin-releasing peptide receptor (GRPR), is also implicated in intrinsic clock network coordination (42). Coupled with the fact that – as in flies – sleep-clock loci do not exhibit mature circuit connectivity in young rodents (43), both the circuit and molecular findings described here might be evolutionarily conserved.

We hypothesize that in immature stages of life, metabolic demand is high and nutritional storage capabilities low, necessitating rapid alterations between sleep and feeding. At this developmental time, clock control of sleep and resultant consolidated sleep periods would be disadvantageous and risk malnourishment. With growth, as prolonged periods without feeding become more sustainable, circadian sleep emerges through a newly-formed clock-to-arousal neuronal circuit. In flies, this rhythmic sleep is deeper at night, unlocking more sophisticated brain functions. Maturation of sleep patterns in human infancy likewise occurs coincidentally with enhanced cognitive capabilities (44–46). Our data suggest a direct link between these events, with onset of daily sleep rhythms enabling persistent memories.

## Acknowledgements

We thank members of the Kayser Lab, Raizen Lab, and members of the Penn Chronobiology and Sleep Institute for helpful discussions/input.

## Author contributions

Conceptualization, A.R.P., A.S.T., D.J.C., M.S.K.; Investigation, A.R.P., L.Z., P.D.M., M.S., R.C.A. Writing – Original Draft, A.R.P. and M.S.K. Writing – Review & Editing, all authors.

## Methods

### Fly Stocks

The following lines have been maintained as lab stocks or were obtained from Dr. Amita Sehgal: iso31, clock^Jrk^ (40), per0 (14), tim0 (13), Dh44^VT^-Gal4 (VT039046) (47), cry-Gal4 pdf-Gal80 (48), UAS-TrpA1 (15), UAS-Kir2.1 (16), elav-Gal4 UAS-dcr2, UAS-mCherry RNAi, VDRC KK control (60100), tsh-Gal80 UAS-TrpA1, tsh-Gal80 UAS-mCD8::GFP, UAS-GCaMP7f UAS-tdTom (49), LexAOP-sp-GFP11; UAS-CD4-spGFP1-10::nrxn (21), pdf-Gal4 (50), LexAOP-GCaMP6 UAS-P2X2 (26). Dh44-LexA (80703), per-Gal4 (7127), cry-Gal4 (24774), CCHa1-R RNAi (51168), CNMaR RNAi (57859), Gyc76D RNAi (57315), pdfr RNAi (38347), Dh44-R1 RNAi (28780), and CCHa1 RNAi (57562) were from the Bloomington Drosophila Stock Center (BDSC). Dh44 KK RNAi (108473) was from the Vienna *Drosophila* Resource Center (VDRC). tub>Gal80>; tsh-LexA LexAOP-Flp; UAS-TrpA1 and tub>Gal80>; tsh-LexA LexAOP-Flp; UAS-GFP were gifts from W. Grueber.

The following lines were used in Fig. S2a (initial neural activation screen) and are maintained as lab stocks: Dh31-Gal4, Akh-Gal4, amn28a-Gal4, AstA-Gal4, hugin-Gal4, NPF-Gal4, Tk-Gal4, ptth-Gal4, Proc-Gal4, Capa-Gal4, Crz-Gal4, CCAP-Gal4, dilp2-Gal4, Dsk-Gal4, Eh-Gal4, ETH-Gal4, FMRFa-Gal4, Lk-Gal4, Ms-Gal4, sNPF-Gal4, Burs-Gal4, pdf-Gal4, Lk-Gal4, SIFa-Gla4, C929-Gal4.

### Larval rearing & sleep assays

Adult flies were maintained on standard molasses-based diet (8.0% molasses, 0.55% agar, 0.2% Tegosept, 0.5% propionic acid) at 25°C on a 12:12 light:dark (LD) cycle. For experiments performed in the subjective evening (CT12-21), adult flies were maintained on a 12:12 reverse dark:light (DL) cycle. In order to collect synchronized second and third instar larvae, adult flies were placed in an embryo collection cage (Genesee Scientific, cat#: 59-100) and eggs were laid on a petri dish containing 3% agar, 2% sucrose, and 2.5% apple juice with yeast paste on top (for 2^nd^ instars) or a molasses-based diet with yeast paste on top (for 3^rd^ instars). Animals developed on this media for two days (for 2^nd^ instars) or three days (for 3^rd^ instars). To examine sleep in either 2^nd^ or 3^rd^ instar larvae at constant conditions, petri dishes were moved to constant darkness (DD) after the first day of entrainment.

Sleep assays for 2^nd^ instar larvae were performed using the LarvaLodge and image acquisition parameters as described previously (5). Briefly, molting 2^nd^ instar larvae were placed into individual wells of the LarvaLodge containing 120 ul of 3% agar and 2% sucrose media covered with a thin layer of yeast paste. Sleep assays in molting 3^rd^ instar larvae were performed using a modified LarvaLodge, with 20 rounded wells of 20 mm diameter & 1.5 mm depth. Molting 3^rd^ instar larvae were placed into individual wells of the modified LarvaLodge containing 95 ul of 3% agar and 2% sucrose media covered with a thin layer of yeast paste. The LarvaLodge was covered with a transparent acrylic sheet and placed into a DigiTherm (Tritech Research) incubator at 25°C for imaging. Experiments were performed in the dark. For thermogenetic experiments, adult flies were maintained at 22°C. Larvae were then placed into the LarvaLodge (as described above) which was moved into a DigiTherm (Tritech Research) incubator at 30°C for imaging.

For analysis of L3 sleep across development while holding circadian time constant, adult flies were maintained in 2 separate incubators on a 12:12 light:dark (LD) cycle set 4 hours apart. Newly molted 3^rd^ instar larvae were selected 1 hour prior to the desired circadian time (for +0 timepoints) and placed into individual wells of the LarvaLodge for sleep analysis. For the +4 and +8 timepoints, newly molted 3^rd^ instar larvae were selected 4 or 8 hours prior to the desired circadian time and then allowed to develop on a petri dish containing a standard molasses-based diet with yeast paste in DD for either 4 hours (+4) or 8 hours (+8). Larvae were then placed into individual wells of the LarvaLodge and experiments were performed as described above.

For analysis of L3 sleep as both circadian time and developmental age proceed across the day, adult flies were maintained in a single incubator on a 12:12 light:dark (LD) cycle. Newly molted 3^rd^ instar larvae were selected and placed into individual wells of the LarvaLodge for sleep analysis (for the first timepoint) or allowed to develop on a petri dish containing a standard molasses-based diet with yeast paste in DD for either 4 hours or 8 hours (for the later timepoints). Larvae were then placed into individual wells of the LarvaLodge and experiments were performed as described above.

For analysis of sleep in animals raised in constant light (LL), larvae on petri dishes in egg lay chambers were entrained to a 12:12 light:dark (LD) or a reverse 12:12 dark:light (DL) cycle for 24 hours. Then the petri dishes were moved to constant light conditions for 48 hours. Molting 3^rd^ instar larvae were moved to dark for adaptation for 3 hours before placing larvae into individual wells of the adapted LarvaLodge. Sleep assays were then conducted in constant darkness.

### LarvaLodge Image acquisition & Processing

Images were acquired every 6 seconds with an Imaging Source DMK 23GP031 camera (2592 × 1944 pixels, The Imaging Source, USA) equipped with a Fujinon lens (HF12.55A-1, 1:1.4/12.5 mm, Fujifilm Corp., Japan) with a Hoya 49mm R72 Infrared Filter. We used IC Capture (The Imaging Source) to acquire time-lapse images. All experiments were carried out in the dark using infrared LED strips (Ledlightsworld LTD, 850 nm wavelength) positioned below the LarvaLodge.

Images were analyzed using custom-written MATLAB software (see Churgin et al 2019 (51) and Szuperak et al 2018 (5)). Temporally adjacent images were subtracted to generate maps of pixel value intensity change. A binary threshold was set such that individual pixel intensity changes that fell below 40 gray-scale units within each well were set equal to zero (“no change”) to eliminate noise. For 3^rd^ instars, the threshold was set to 45 to account for larger body size. Pixel changes greater than or equal to threshold value were set equal to one (“change”). Activity was then calculated by taking the sum of all pixels changed between images. Sleep was defined as an activity value of zero between frames. For 2^nd^ instar sleep experiments done across the day, total sleep was summed over 6 hrs beginning 2 hrs after the molt to second instar (Fig. 2 & Fig. S2). For sleep experiments performed at certain circadian times, total sleep in the 2^nd^ hour after the molt to second (or third) instar was summed. For all experiments, sleep metrics were normalized to the average value for the control for a given biological replicate.

For experiments examining development and circadian interactions (Fig. S1), activity values were normalized to account for increasing body size over the experimental day. We adjusted the binary thresholding values to define sleep episodes as less than 2% change in activity.

### Sleep bout distribution analysis

Sleep bout distribution analysis was performed as described previously (52). Briefly, we calculated the duration of every sleep bout for the 2^nd^ hour of the experiment for L2 & L3 larvae at CT1 and CT13. All sleep bouts for a given condition were pooled together. The pooled data was then used to generate cumulative relative frequency plots. The cumulative relative frequency for any given sleep episode duration (sec) was calculated as the total number of bouts equal to and shorter than that sleep episode duration divided by the total number of all bouts. The resulting cumulative relative frequency was plotted on the y-axis and the x-axis represented the continuous time interval with a bin size of 6 sec. For comparing sleep bout distributions, Kolmogorov-Smirnov tests were used.

### Calculation of p(wake) and p(doze)

p(wake) and p(doze) were calculated as described previously (38). p(doze) is defined as the number of wake (active) to sleep (quiescent) transitions divided by the number of frames awake (active). p(wake) is defined as the number of sleep (quiescent) to wake (active) transitions divided by the number of frames asleep (quiescent).

### Feeding behavior analysis

Newly molted 2^nd^ instar larvae were placed in individual wells of the LarvaLodge containing 120 ul of 3% agar and 2% sucrose media covered with a thin layer of yeast paste. Larvae were then imaged continuously with a Sony HDR-CX405 HD Handycam camera (B&H Photo, Cat. No: SOHDRCX405) for 10 minutes. The number of mouth hook contractions (feeding) was counted manually over the imaging period and divided by the time awake.

### Aversive Olfactory conditioning

We used an established two odor reciprocal olfactory conditioning paradigm with 10 mM quinine (quinine hydrochloride, EMSCO/Fisher, Cat. No: 18-613-007) as a negative reinforcement to test short-term or long-term memory performance in L2 and early L3 larvae (30) at CT12-15. Experiments were conducted on assay plates (100 × 15 mm, Genesee Scientific, Cat. No: 32-107) filled with a thin layer of 2.5% agarose containing either pure agarose (EMSCO/Fisher, Cat. No: 16500-500) or agarose plus reinforcer. As olfactory stimuli, we used 10 μl amyl acetate (AM, Sigma-Aldrich, Cat. No: STBF2370V, diluted 1:50 in paraffin oil-Sigma-Aldrich, Cat. No: SZBF140V) and octanol (OCT, Fisher Scientific, Cat. No: SALP564726, undiluted). Odorants were loaded into the caps of 0.6 mL tubes (EMSCO/Fisher, Cat. No: 05-408-123) and covered with parafilm (EMSCO/Fisher, Cat. No: 1337412). Larvae were trained by exposing a group of 30 larvae to AM while crawling on agarose medium plus quinine reinforcer. After 5 min, larvae were transferred to a fresh Petri dish containing agarose alone with OCT as an odorant (AM+/OCT). A second group of 30 larvae received the reciprocal training (AM/OCT+). Three training cycles were used for all experiments. For long-term memory, larvae were transferred after training onto agarose plates with a small piece of Kimwipe moistened with tap water and covered in dry active yeast (LabScientific, Cat. No: FLY804020F). Larvae were then kept in the dark for 1.5 hrs before testing memory performance. For short-term memory, larvae were immediately transferred after training onto test plates (agarose plus reinforcer) on which AM and OCT were presented on opposite sides of the plate. After 5 min, individuals were counted on the AM side, the OCT side, or in the middle. We then calculated a preference index (PREF) for each training group by subtracting the number of larvae on the unconditioned stimulus side from the number of larvae on the conditioned stimulus side. We then took the average of each PREF value to calculate an associative performance index (PI) as a measure of associative learning.

### Arousal and sleep deprivation

Blue light stimulation was delivered as described in (5) using 2 high power LEDs (Luminus Phatlight PT-121, 460 nm peak wavelength, Sunnyvale, CA) secured to an aluminum heat sink. The LEDs were driven at a current of 0.1 A (low intensity) (for arousal experiments) or 1 A (high intensity) (for sleep deprivation experiments). For sleep deprivation, we used a high intensity stimulus for 30 sec every 2 minutes for 1 hr beginning the 2^nd^ hr after the molt to third instar. Undisturbed control animals were placed in a separate incubator during deprivation. For arousal experiments, we used a low intensity stimulus for 4 sec every 2 minutes for 1 hr beginning the 2^nd^ hr after the molt to second (or third) instar. We then counted the number of larvae that showed an activity change in response to stimulus. To compare arousal thresholds in 2^nd^ instar larvae to those in larger 3^rd^ instar larvae at different CT times, we normalized the activity values for 3^rd^ instars to their relative body size. We used the standard binary thresholding to define sleep episodes and then subtracted 35 pixels to achieve a similar degree of change in response to the light stimulus.

### Immunohistochemistry & imaging

Brains were dissected in PBS, fixed in 4% PFA for 20 min at room temperature. Following 3 × 20 min washes in PBST, brains were incubated with primary antibody at 4°C overnight. Following 3 × 20 min washes in PBST, brains were incubated with secondary antibody at 4°C overnight. Following 3 × 20 min washes in PBST, brains were mounted in Vectashield (Vector Laboratories. Primary antibodies included: Rabbit anti-GFP (1:200, A-11122, ThermoFisher Scientific), Mouse anti-GFP (for GRASP dissections) (1:200, EMSCO/Fisher, Cat. No: G6539-.2ML), and Rabbit anti-Dh44 (1:500, Johnson et al 2005). Secondary antibodies included Alexa Fluor donkey antirabbit 488 (1:500, Jackson) and goat anti-mouse 488 (1:500, Jackson). Brains were visualized and imaged with a Leica SP8 confocal microscope.

### P2X2 Activation and GCaMP imaging

For all live imaging experiments (GCaMP, P2X2, and CCHa1 bath application), brains were dissected in AHL buffer consisting of (in mM): 108 NaCl, 5 KCl, 2 CaCl2, 8.2 MgCl2, 4 NaHCO3, 1 NaH2PO4-H20, 5 Trehalose, 10 Sucrose, 5 HEPES, pH=7.5. Brains were placed on a small glass coverslip (Carolina Cover Glasses, Circles, 12 mm, Cat. No: 633029) in a perfusion chamber filled with AHL. Solutions were perfused over the brains using a gravity-fed ValveLink perfusion system.

For baseline GCaMP7f imaging at CT0 and CT12, brains were chosen for imaging based on UAS-tdTomato signal. AHL buffer was perfused over the brains throughout imaging. Twelve-bit images were acquired with a 40 × water immersion objective at 256 × 256-pixel resolution. Z-stacks were acquired every 30 sec for 10 min. Image processing and measurement of fluorescence intensity was performed in ImageJ. A max intensity Z-projection of each time step and Smooth thresholding was used for analysis. Regions of interest (ROIs) were manually drawn in ImageJ to encompass individual tdTomato-positive cell bodies and mean fluorescence intensities were measured for each ROI at each time point. GCaMP7f signal was normalized to tdTomato signal (FGCaMP7f/tdTom) for each ROI. For each cell, the average FGCaMP7f/tdTom over the 10 min imaging period was used as a measure of relative fluorescence (A.U.). All analysis was done blind to experimental condition.

For P2X2 imaging, dissections were performed at CT12-15 and AHL buffer was perfused over the brains for 1 min of baseline GCaMP6 imaging, then ATP was delivered to the chamber by switching the perfusion flow from the channel containing AHL to the channel containing 2.5 mM ATP in AHL, pH 7.5. ATP was perfused for 2 min. Twelve-bit images were acquired with a 40 × water immersion objective at 256 × 256-pixel resolution. Z-stacks were acquired every 5 sec for 3 min. Image processing and measurement of fluorescence intensity was performed in ImageJ. A max intensity Z-projection of each time step and Smooth thresholding was used for analysis. Regions of interest (ROIs) were manually drawn in ImageJ to encompass individual GCaMP-positive cell bodies and mean fluorescence intensities were measured for each ROI at each time point. For each cell body, fluorescence traces over time were normalized using this equation: ΔF/F = (F_n_-F_0_)/F_0_, where F_n_=fluorescence intensity recorded at time point n, and F_0_ is the average fluorescence value during the 1 min baseline recording. All analysis was done blind to experimental condition.

For CCHa1 bath application, dissections were performed at CT12-15 and AHL buffer was perfused over the brains for 1 min of baseline GCaMP7f imaging, then CCHa1 peptide was delivered to the chamber by switching the perfusion flow from the channel containing AHL to the channel containing 1 μM synthetic CCHa1 in AHL, pH 7.5. CCHa1 was perfused for 2 min, followed by a 1 min wash-out with AHL. For the AHL negative control, the perfusion flow was switched from one channel containing AHL to another channel containing AHL. Twelve-bit images were acquired with a 40 × water immersion objective at 256 × 256-pixel resolution. Z-stacks were acquired every 10 sec for 4 min. Image processing and measurement of fluorescence intensity was performed in ImageJ. A max intensity Z-projection of each time step and Smooth thresholding was used for analysis. Image analysis was performed in a similar manner as for the P2X2 experiments. All analysis was done blind to experimental condition.

### Statistical analysis

All statistical analysis was done in GraphPad (Prism). For comparisons between 2 conditions, two-tailed unpaired *t*-tests were used. For comparisons between multiple groups, ordinary one-way ANOVAs followed by Tukey’s multiple comparison tests were used. For comparisons between different groups in the same analysis, ordinary oneway ANOVAs followed by Sidak’s multiple comparisons tests were used. For comparison of p(wake) and p(doze) values, a two-sided Wilcoxon Rank-Sum test was used. For comparison of GCaMP signal in P2X2 and CCHa1 experiments, Mann-Whitney test was used. For comparison of sleep bout distributions, Kolmogorov-Smirnov tests were used. \**P*<0.05, \*\**P*<0.01, \*\*\**P*<0.001. Representative confocal images are shown from at least 8-10 independent samples examined in each case. For live imaging, ~15-20 neurons in 5-10 brains were used for each condition.

---

**Fig. S1:**
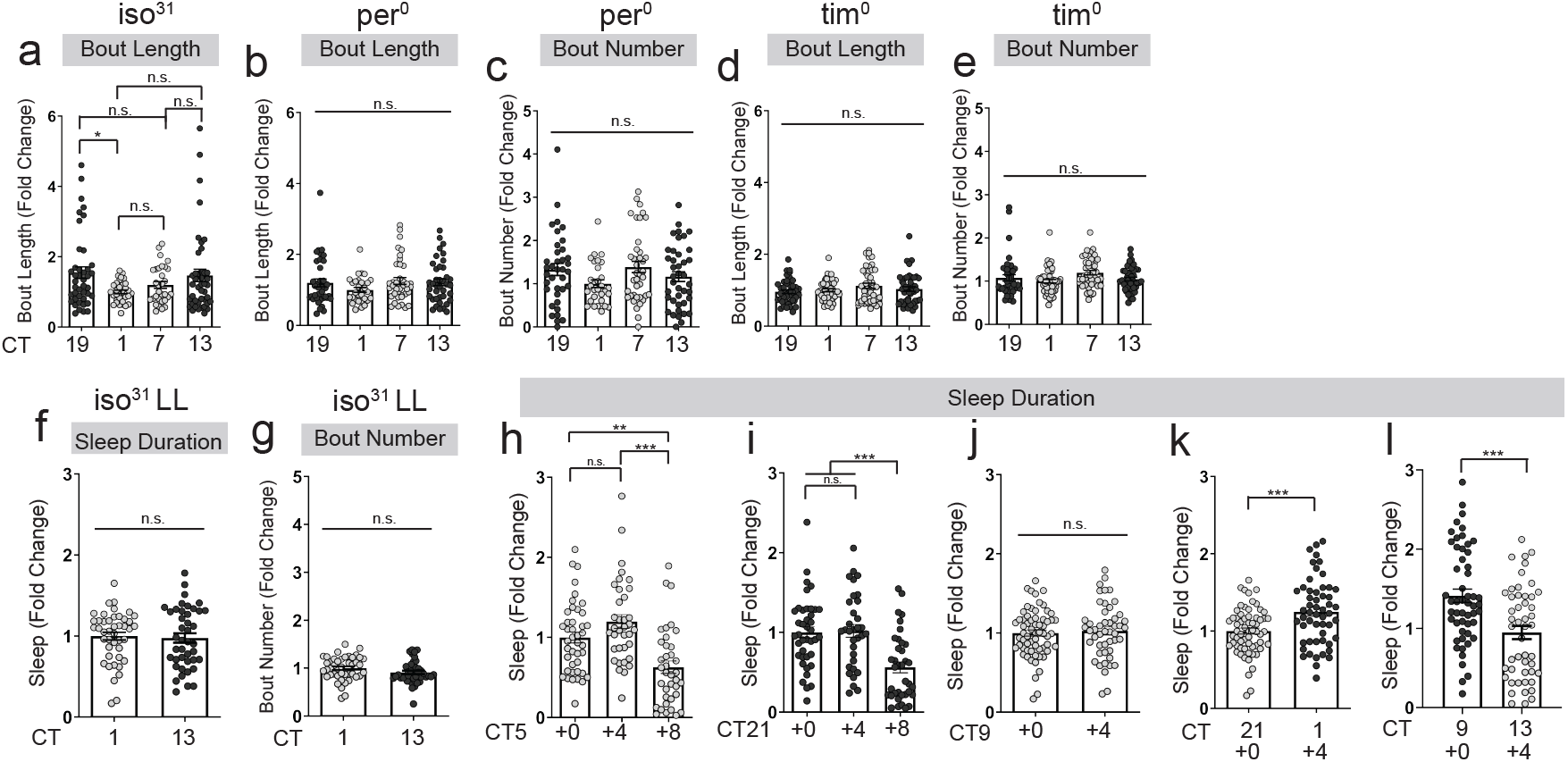
Early L3 show rhythmic differences in sleep. **a**, Bout length of L3 controls across the day. **b-e**, Sleep bout length and number of L3 clock mutants. **f-g**, Sleep duration and bout number in L3 reared in constant light (LL). **h-j**, Sleep duration at CT5 (h), CT21 (i), and CT9 (j) in newly molted L3 (+0), larvae aged 4 hrs (+4), and larvae aged 8 hrs (+8). **k-l**, Sleep duration in developing L3 larvae during the CT21 to CT1 transition (k) and CT9 to CT13 transition (l). Subjective day is shown in light gray circles, subjective night in black. All sleep metrics represent fold change (normalized to avg. value of control). a-e, n=31-44 larvae; f-g, n=42-44 larvae; h-l, n=40-60 larvae. One-way ANOVA followed by Tukey’s multiple comparison test [(a-e) and (h-i)] and unpaired twotailed Student’s *t*-test [(f-g) and (j-l)]. n.s., not significant, \**P*<0.05, \*\**P*<0.01, \*\*\**P*<0.001 in this and all subsequent supplemental figures.

**Fig. S2:**
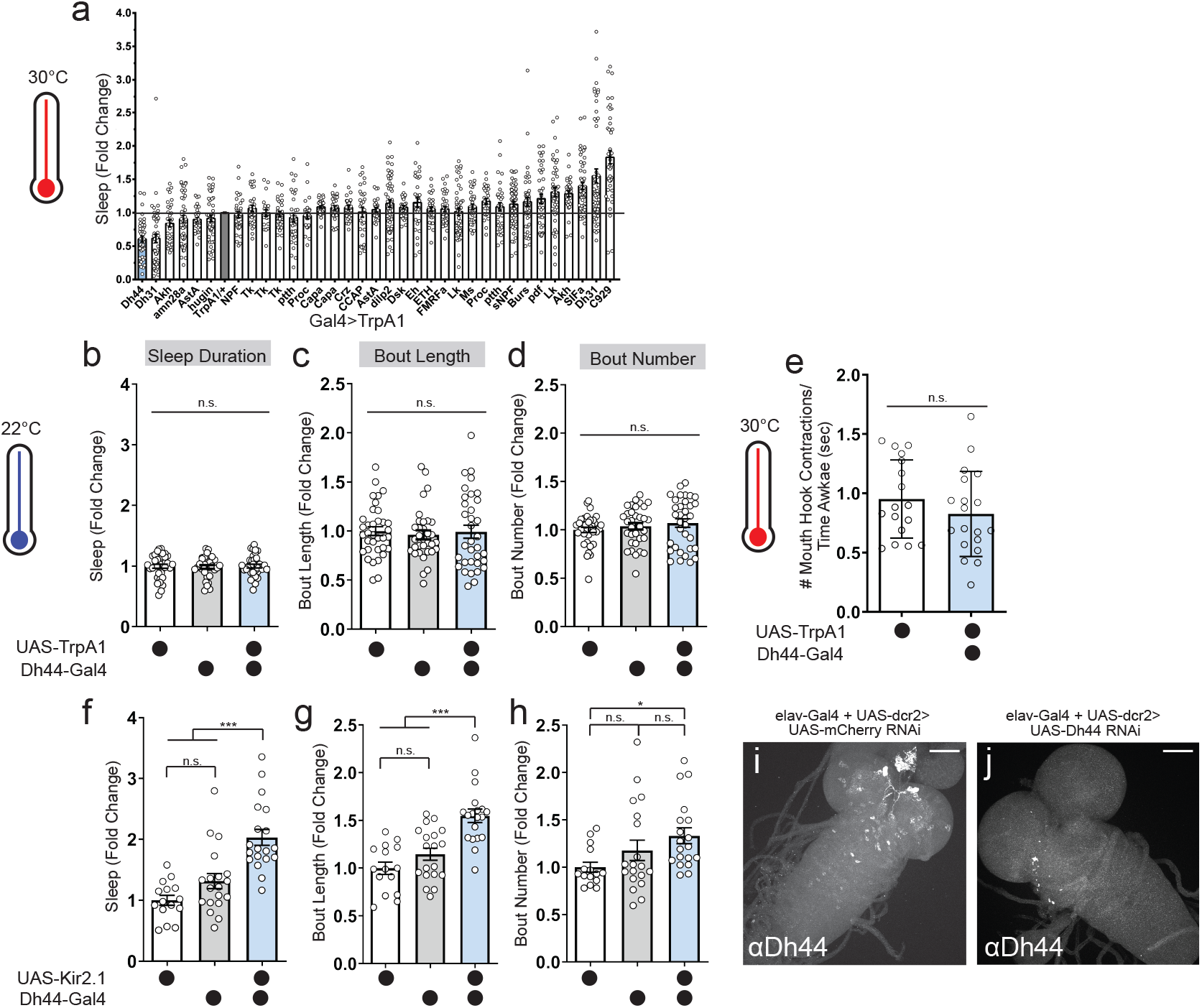
Dh44 modulates L2 waking. **a**, Sleep duration of neuropeptidergic activation screen (*Gal4*>*UAS-TrpA1*) in L2 at 30°C. **b-d**, Sleep duration, bout length, and bout number in L2 expressing *UAS-TrpA1* with *Dh44-Gal4* and genetic controls at 22°C. **e**, Normalized feeding (# mouth hook contractions/time awake (sec)) in L2 expressing *UAS-TrpA1* with *Dh44-Gal4* and controls at 30°C. **f-h**, Sleep duration, bout length, and bout number in L2 expressing *UAS-Kir2.1* with *Dh44-Gal4* and genetic controls. **i-j**, Images of *elav-Gal4*>*UAS-dcr2* + *UAS-mCherry RNAi* (j) and *elav-Gal4*>*UAS-dcr2* + *UAS-Dh44 RNAi* (k) L2 brains immunostained for Dh44. n=5 brains. All sleep metrics represent fold change (normalized to avg. value of control). a, n=30-50 larvae; b-d, n=30 larvae; e, n=18-20 larvae; f-h, n=14-18 larvae. One-way ANOVA followed by Tukey’s multiple comparison test [(b-d) and (f-h)] and unpaired two-tailed Student’s *t*-test (e). Scale bars=50 microns.

**Fig. S3:**
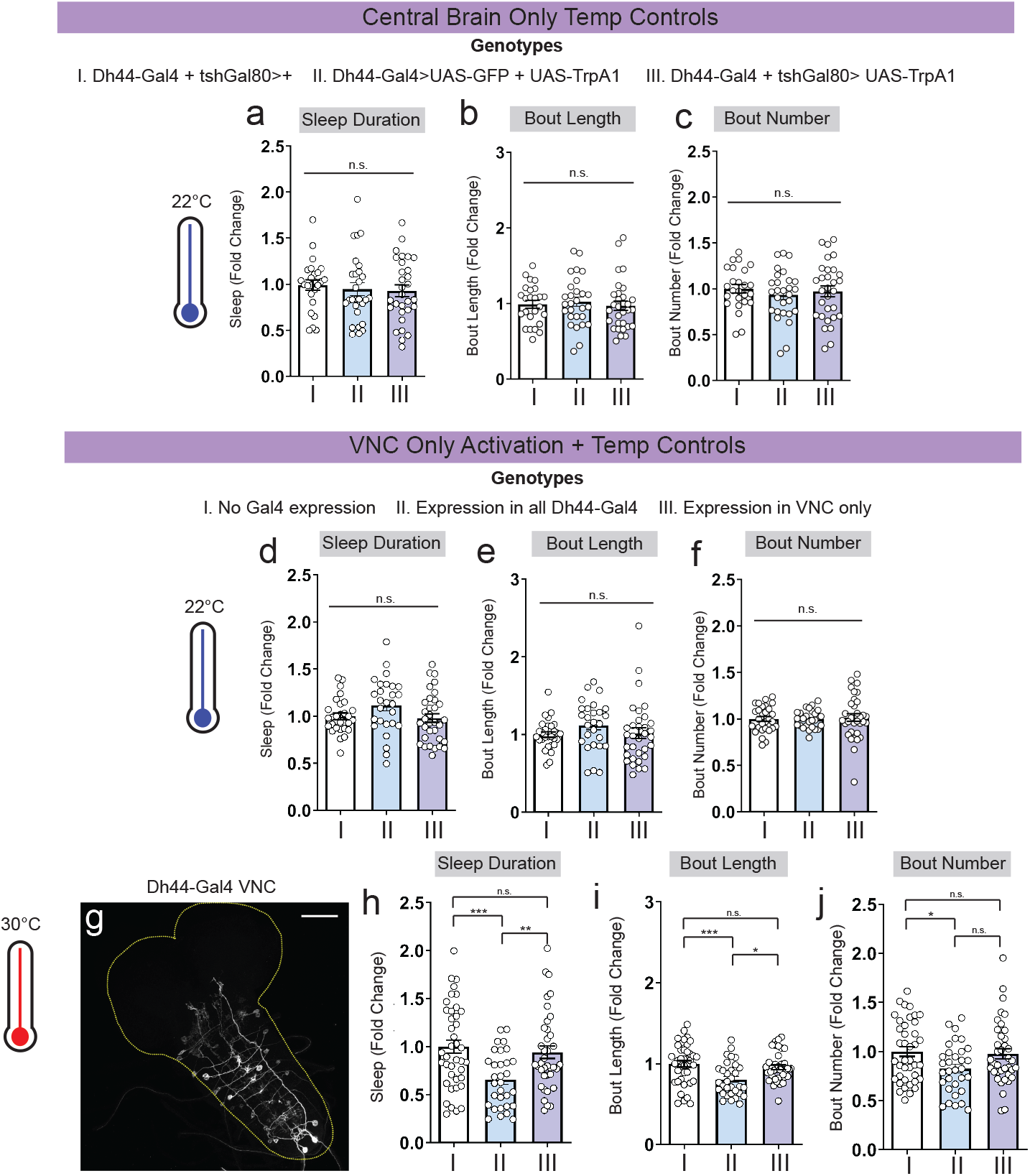
Intersectional analysis of Dh44 neurons in L2. **a-c**, Sleep duration, bout length, and bout number in L2 expressing *Dh44*-Gal4 in presence of *tshGal80* and *UAS-TrpA1* with genetic controls at 22°C. **d-f**, Sleep duration, bout length, and bout number of VNC-only Dh44 neurons and genetic controls at 22°C. **g**, L2 brain and ventral nerve cord showing GFP expression in larvae expressing *Dh44*-Gal4 only in VNC. **h-j**, Sleep duration, bout length, and bout number with thermogenetic activation of VNC-only Dh44 neurons and genetic controls at 30°C. All sleep metrics represent fold change (normalized to avg. value of control). a-c, d-f, n=25-30 larvae; h-j, n=30-40 larvae. Oneway ANOVA followed by Tukey’s multiple comparison test. Scale bars=50 microns.

**Fig. S4:**
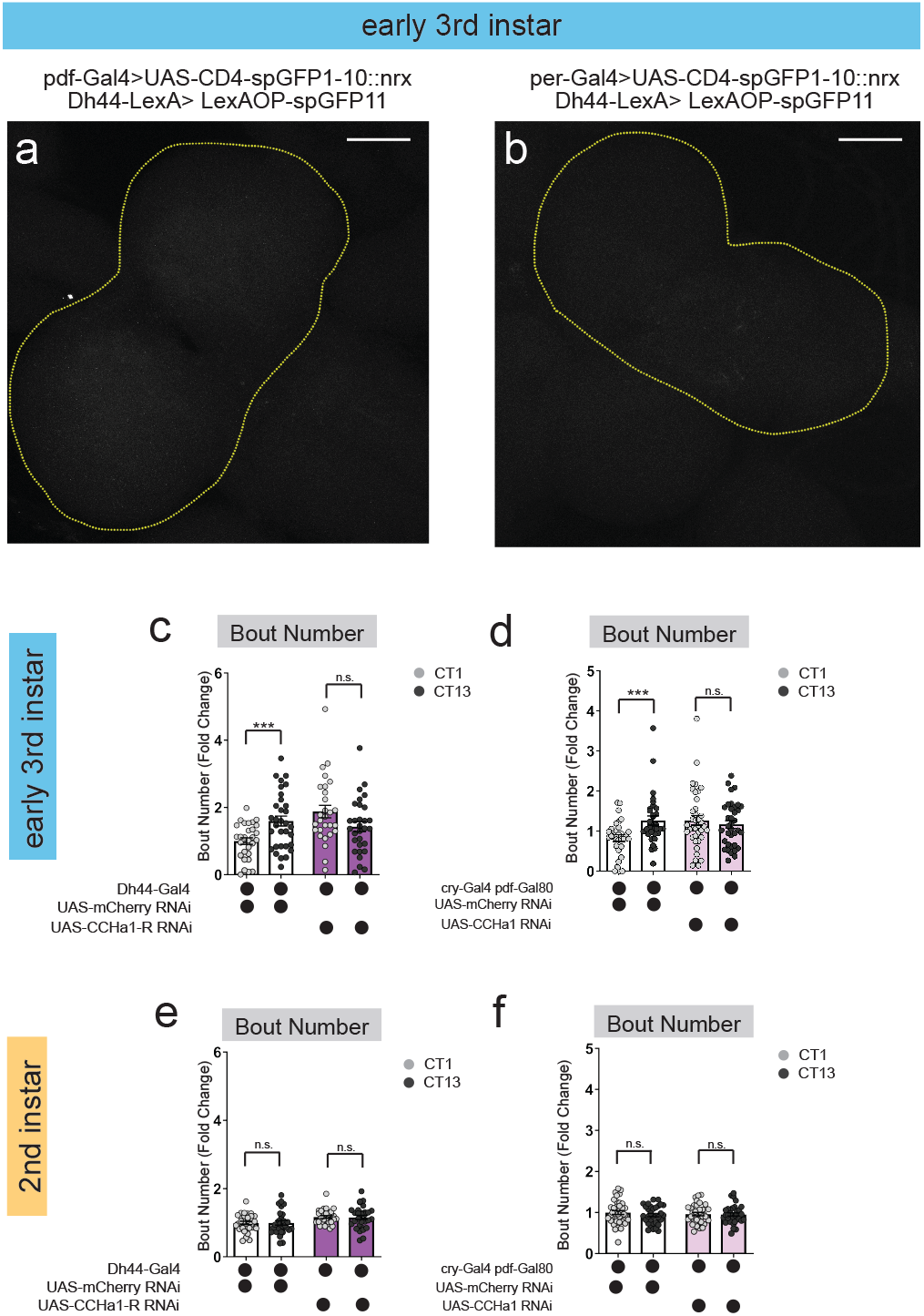
DN1as regulate sleep rhythms in early L3. **a-b**, Absence of neurexin-based GFP reconstitution (GRASP) between s-LNvs (*pdf*-Gal4) (a) or DN2s (*per*-Gal4) (b) and Dh44 neurons (*Dh44*-LexA) in L3 brains. Yellow dotted lines indicate central brain region. n=8-10 brains. **c-d**, Sleep bout number in L3 expressing *UAS-CCHa1-R-RNAi* with *Dh44-Gal4* (c) or *UAS-CCHa1-RNAi* with *cry-Gal4 pdf-Gal80* (DN1as) (d) and genetic controls at CT1 and CT13. **e-f**, Sleep bout number in L2 expressing *UAS-CCHa1-R-RNAi* with *Dh44-Gal4* (e) or *UAS-CCHa1-RNAi* with *cry-Gal4 pdf-Gal80* (DN1as) (f) and genetic controls at CT1 and CT13. Subjective day is shown in light gray circles, subjective night in black. All sleep metrics represent fold change (normalized to avg. value of control). c-f, n=29-38 larvae. Ordinary one-way ANOVAs followed by Sidak’s multiple comparisons tests. Scale bars=50 microns.

**Fig. S5:**
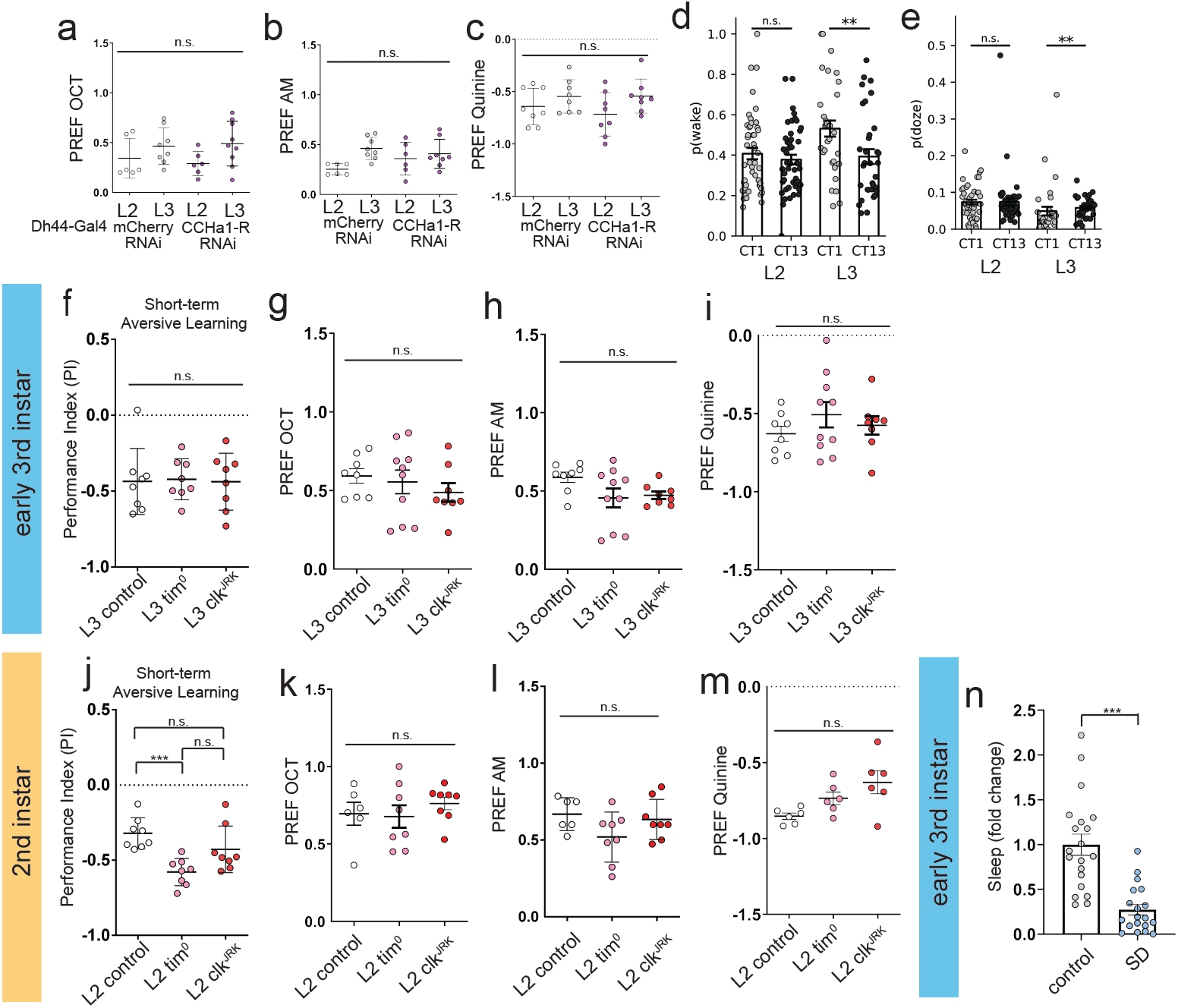
Naïve preferences for odor and aversive stimuli in L2 and L3. **a-c**, Naïve OCT, AM, and quinine preference in L2 and L3 expressing *Dh44-Gal4*>*mCherry RNAi* and *Dh44-Gal4*>*CCHa1-R-RNAi*. **d-e**, Probability of transitioning from a wake state to sleep [p(wake)] (d) and probability of exiting a sleep state [p(doze)] (e) in L2 and L3 at CT1 and CT13. **f**, Short-term aversive memory performance in L3 controls, *tim^0^*, and *clk^JRK^* mutants. **g-i**, Naïve OCT, AM, and quinine preference in L3 controls, *tim^0^*, and *clk^JRK^* mutants. **j**, Short-term aversive memory performance in L2 controls, *tim^0^*, and *clk^JRK^* mutants. **k-m**, Naïve OCT, AM, and quinine preference in L2 controls, *tim^0^*, and *clk^JRK^* mutants. **m**, Quantification of sleep loss during sleep deprivation (SD) over 1 hr (fold change). a-c, f-h, k-m, n=6 PREFs (180 larvae) per genotype; i-j, n= n=8 Pls (240 larvae) per genotype; n, n=20-22 larvae. One-way ANOVA followed by Tukey’s multiple comparison test [(a-c), (f-m)] plus two-sided Wilcoxon Rank-Sum test [(d-e)] and unpaired two-tailed Student’s *t*-test (n).

